# Static allometry of the horn and pronotal depression in adult *Oryctes rhinoceros* supports continuous nonlinearity, sexual dimorphism, and size-independent covariation

**DOI:** 10.64898/2026.05.28.728611

**Authors:** Manato Okahara, Teruyuki Niimi, Shinichi Morita

## Abstract

Exaggerated insect traits often show positive allometry, yet nonlinear scaling can reflect either continuous curvature or discrete morphs. Distinguishing between these alternatives is important because they imply different developmental and evolutionary scenarios. Using cross-sectional data from 1,000 adult *Oryctes rhinoceros*, we analyzed the static allometry of nine traits with pronotum width as the primary body-size proxy and body length for sensitivity analyses. Cross-validated comparisons among linear, continuous nonlinear, and two-component mixture models showed that continuous nonlinear models improved predictive performance over linear models for horn length and pronotal depression width in both sexes. By contrast, mixture regression did not outperform the best continuous model in either sex, providing no positive support for discrete within-sex dimorphism under the tested model set. After accounting for body size, horn length remained positively associated with pronotal depression width, indicating size-independent covariation. These associations were retained when body length was used instead of pronotum width, supporting robustness to body-size proxy choice. Together, these results support continuous nonlinear adult scaling of the horn and pronotal depression in *O. rhinoceros* and indicate covariation not attributable solely to body size under the tested model set.

## Introduction

Exaggerated traits such as weapons and sexual ornaments often mediate intraspecific competition and mating success^1,2^. In adult populations, these structures frequently exhibit positive allometry, increasing disproportionately with body size^2,3^. Yet nonlinear static allometry can arise in at least two conceptually different ways: as continuous curvature or as discrete dimorphism associated with a body-size threshold^4^. Distinguishing between these alternatives matters because they imply different developmental mechanisms and different interpretations of the adaptive significance of exaggerated traits^5^. Moreover, body-size thresholds can evolve rapidly^6^, making this distinction relevant to both developmental and evolutionary analyses. In horned beetles, exaggerated structures arise primarily on the head and pronotum, the principal regions of horn formation^7^.

The coconut rhinoceros beetle, *Oryctes rhinoceros* (Coleoptera: Scarabaeidae), is a major pest of palms^8^. Because females as well as males bear a head horn, this species provides an informative system for examining sexual dimorphism and the functional evolution of exaggerated traits. However, the allometric scaling of the horn has not yet been quantified in both sexes of this species. In addition to the head horn, *O. rhinoceros* possesses a deep broad depression on the pronotum^9^, here termed the pronotal depression. It has likewise not been tested quantitatively how these traits scale with body size or whether their covariation persists after body size has been taken into account.

Our first objective was to test, on the basis of generalization performance, whether cross-sectional adult scaling patterns were better described by continuous nonlinear models than by linear models, and whether they required an explicit dimorphism model. Using pronotum width (PW) as the primary body-size proxy, we (i) compared the static allometry of nine traits across a candidate set that crossed alternative mean structures (LINEAR, LOGISTIC, SEGMENTED, and GAM) with homoscedastic and heteroscedastic variance structures; (ii) quantified sexual dimorphism for the focal nonlinear traits by comparing fitted curves and predicted values at representative PW values; and (iii) evaluated the discrete-dimorphism hypothesis by comparing two-component mixture regression (MIXTURE_K2) with the best continuous alternatives. We further (iv) tested whether horn and pronotal depression covaried beyond their shared dependence on body size; and (v) assessed the robustness of the main conclusions by repeating the analyses with body length (BL) as an alternative body-size proxy. Overall, the horn and pronotal depression were best characterized by continuous nonlinearity and size-independent covariation rather than by discrete within-sex dimorphism under the tested model set. These analyses provide a quantitative framework for understanding the static allometry of exaggerated and related traits in *O. rhinoceros*.

## Materials and Methods

### Oryctes rhinoceros

Adult *Oryctes rhinoceros* were collected in pheromone traps on Minami-Daito Island, Okinawa, Japan, as part of an invasive-species control program conducted under the Okinawa Prefecture action plan for invasive alien species. Specimens were kindly provided by Minami-Daito Village Office and Minami-Daito Village Agricultural Youth Club. Some individuals had lost tarsal segments or other appendages during collection and transport, but only specimens in which such damage did not affect measurement of the nine focal traits were retained for analysis. Specimens were selected to span the body-size range broadly in both sexes while avoiding strong bias toward any particular size class, rather than to reproduce the natural size-frequency distribution. Accordingly, the dataset was designed for cross-sectional curve estimation and was not used to estimate morph frequencies or population size structure. The final dataset comprised 500 males and 500 females. Sex was determined from the presence or absence of terminal abdominal hairs^10^ and from the morphology of the abdominal sternites (Fig. S1).

### Trait definitions and measurements

Pronotum width (PW) served as the primary proxy for overall body size. We analyzed nine traits: horn length (HL), pronotal depression length (PDL), pronotal depression width (PDW), body length (BL), elytral length (EL), metafemur length (ML), pronotum length (PL), abdominal width (AW), and abdominal depth (AD). By comparing the exaggerated trait, the related pronotal trait, and additional body traits within a common analytical framework, we assessed whether strong nonlinearity was confined to particular traits. Measurement positions for all traits are shown in Fig. 1. Because PDL and PDW are potentially sensitive to how the boundary of the depression is delineated, landmarks defining the contour were specified a priori to standardize measurements (Fig. S2). To assess reproducibility for PDL and PDW, 50 males and 50 females were randomly selected for remeasurement (Table S1). Reproducibility was quantified using the two-way absolute-agreement single-measure intraclass correlation coefficient (ICC)^11^, Lin’s concordance correlation coefficient (CCC)^12^, the technical error of measurement (TEM)^13^, and relative TEM (%). All measurements were taken by a single observer using digital calipers (DN-100, Niigata Seiki Co., Ltd., Japan).

**Fig. 1.**
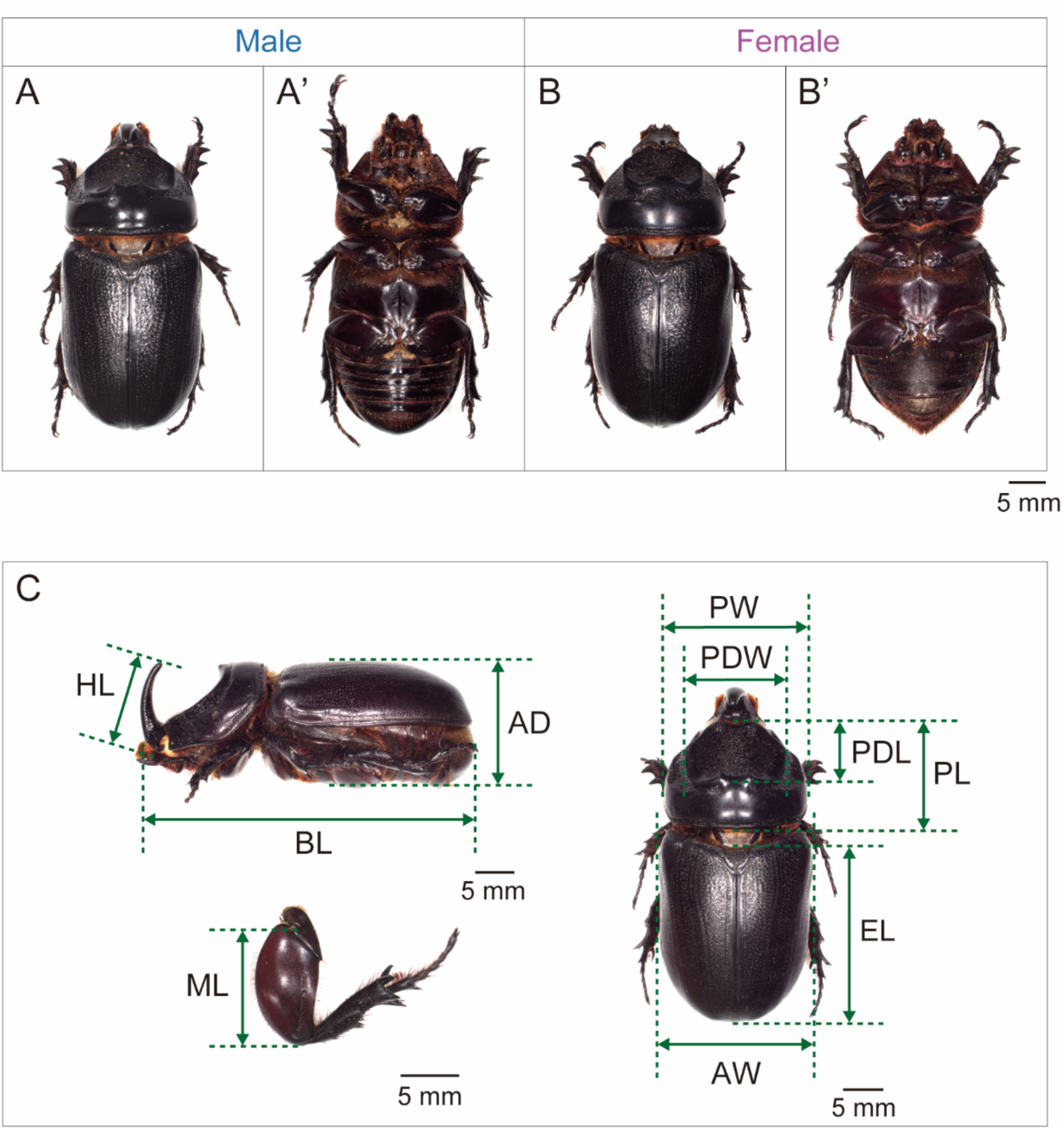
Adult males and females of *O. rhinoceros* and definitions of the measured traits. Dorsal (A) and ventral (A’) views of an adult male, dorsal (B) and ventral (B’) views of an adult female. (C) Measurement sites for the ten traits including PW. PW, pronotum width; HL, horn length; PDL, pronotal depression length; PDW, pronotal depression width; BL, body length; EL, elytral length; ML, metafemur length; PL, pronotum length; AW, abdominal width; AD, abdominal depth.

To facilitate visual comparison of within-sex and between-sex variation in trait size across PW, representative individuals were selected at 1-mm PW intervals from 13 to 20 mm for each measured trait (Fig. S3). Images were acquired with a digital microscope (VHX-900, KEYENCE Co., Japan).

### Statistical analyses

All analyses were conducted in R version 4.4.2. Generalized additive models (GAMs) were fitted with mgcv^14^. Piecewise linear models (SEGMENTED) and two-component mixture regression (MIXTURE_K2) were implemented as custom functions based on maximum-likelihood estimation with optim and on the EM algorithm, respectively, following standard approaches to segmented regression^15^ and mixture regression^16^.

To model the relationship between PW and each trait, we considered four alternative mean structures: a linear model (LINEAR), a four-parameter saturating logistic model (LOGISTIC), a one-breakpoint piecewise linear model (SEGMENTED), and a generalized additive model (GAM). Each mean model was crossed with either a homoscedastic variance structure (HOM), which assumes constant residual variance, or a heteroscedastic variance structure (HET), in which residual variance varies with PW. We therefore compared eight continuous models: LINEAR_HOM/HET, LOGISTIC_HOM/HET, SEGMENTED_HOM/HET, and GAM_HOM/HET.

For the HET models, we first fitted the mean model and then modeled the squared residuals with a Gamma generalized linear model with a log link, using PW and PW^2 as predictors, to estimate the predicted residual variance, sigma^2(PW), as a function of PW^17^. This procedure allowed improvements attributable to mean structure (nonlinearity) and variance structure (heteroscedasticity) to be compared on a common negative log-likelihood (nll) scale. Influential observations were identified using Cook’s distance and studentized residuals (Fig. S4)^18,19^. Because such points were rare within each trait and most appeared as isolated observations in the scatterplots, no exclusion rule was imposed and all specimens were retained in the analyses (Table S2).

Model selection was based on repeated 10-fold cross-validation (20 repeats)^20^. For each candidate model, nll was evaluated on the test data, and the model with the lowest mean nll was taken as the best continuous model for each trait and sex. To avoid unnecessary complexity, we also applied the one-standard-error rule^21^ to the continuous model set (including the linear model) and adopted, as a conservative choice, the simplest model within one standard error of the minimum mean nll (Best-fit model [1SE]) (Table 1; Table S3; Fig. 2; Fig. S5).

**Fig. 2.**
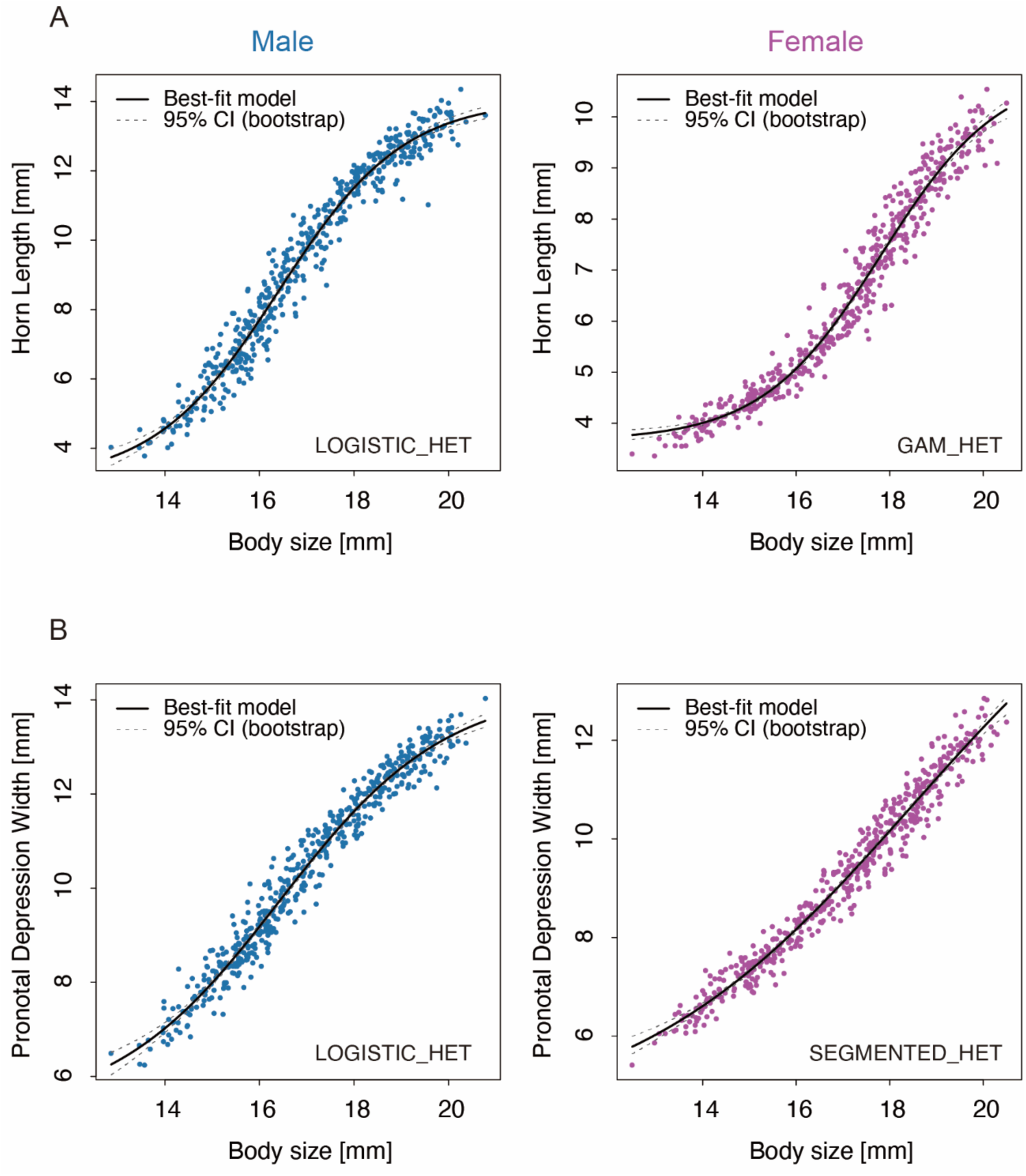
Static allometry of HL and PDW with 95% confidence bands. Fitted curves for HL and PDW against PW are shown with 95% confidence bands. The best-supported model selected by the 1-SE rule (Best-fit model [1SE]) is shown by the solid black line, and the 95% confidence bands are shown by gray dashed lines. Males are shown in blue and females in magenta. (A) HL: LOGISTIC_HET in males and GAM_HET in females. (B) PDW: LOGISTIC_HET in males and SEGMENTED_HET in females. Corresponding curves for the remaining seven traits are shown in Fig. S5.

**Table 1.** Best-fit model [1SE] and the associated Δnll and Δnll_HET values from repeated cross-validation. Δnll = nll (LINEAR_HOM) − nll (Best-fit model [1SE]). Δnll_HET = nll (LINEAR_HET) − nll (Best-fit model [1SE]).

| Trait | Sex | Best-fit model [1SE] | $\Delta nll$ [95%CI] | $\Delta nll\_HET$ |
| --- | --- | --- | --- | --- |
| HL | Male | LOGISTIC_HET | 0.186 [0.184, 0.188] | 0.197 |
| HL | Female | GAM_HET | 0.387 [0.385, 0.389] | 0.395 |
| PDL | Male | LOGISTIC_HOM | 0.053 [0.052, 0.054] | — |
| PDL | Female | LOGISTIC_HET | 0.020 [0.019, 0.020] | 0.025 |
| PDW | Male | LOGISTIC_HET | 0.095 [0.094, 0.096] | 0.105 |
| PDW | Female | SEGMENTED_HET | 0.092 [0.091, 0.093] | 0.094 |
| BL | Male | LINEAR_HET | 0.018 [0.017, 0.019] | 0.003 |
| BL | Female | LINEAR_HET | 0.012 [0.011, 0.013] | 0.004 |
| EL | Male | LOGISTIC_HET | 0.044 [0.043, 0.045] | 0.024 |
| EL | Female | GAM_HOM | 0.006 [0.006, 0.007] | — |
| ML | Male | LOGISTIC_HOM | 0.007 [0.006, 0.008] | — |
| ML | Female | LOGISTIC_HOM | 0.020 [0.020, 0.021] | — |
| PL | Male | LINEAR_HET | 0.008 [0.007, 0.008] | 0.003 |
| PL | Female | SEGMENTED_HET | 0.006 [0.005, 0.008] | 0.005 |
| AW | Male | LINEAR_HET | 0.001 [ $9.2 \times 10^{-5}$, 0.002] | 0.001 |
| AW | Female | LOGISTIC_HOM | 0.004 [0.003, 0.005] | — |
| AD | Male | LOGISTIC_HOM | 0.004 [0.003, 0.005] | — |
| AD | Female | LOGISTIC_HOM | 0.004 [0.004, 0.005] | — |

The magnitude of nonlinear improvement over the linear homoscedastic model was defined as Delta nll = nll (LINEAR_HOM) − nll (Best-fit model [1SE]) (Table 1; Fig. 3; Fig. S6). When the Best-fit model [1SE] was an HET model, we additionally calculated Delta nll_HET = nll (LINEAR_HET) − nll (Best-fit model [1SE]) and compared it with Delta nll to assess whether the improvement was attributable mainly to nonlinearity in the mean structure or to heteroscedasticity in the variance structure (Table 1).

**Fig. 3.**
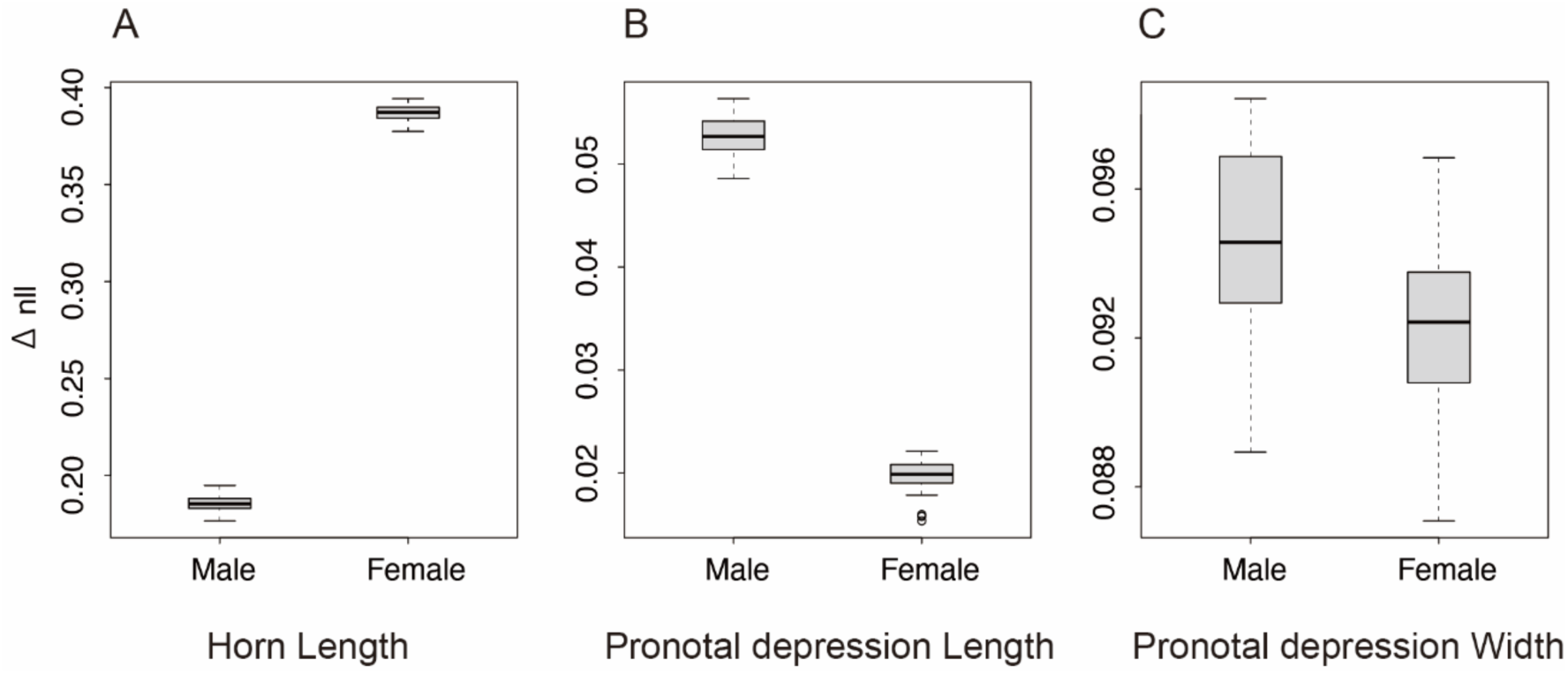
Distribution of Δnll from repeated cross-validation. Boxplots show Δnll = nll (LINEAR_HOM) − nll (Best-fit model [1SE]) from repeated cross-validation (20 repeats) for HL, PDL, and PDW. Larger positive Δnll values indicate better predictive performance of the continuous nonlinear model than of the linear homoscedastic model. Distributions for the other six traits are shown in Fig. S6.

Sex differences in trait size were quantified by fitting the Best-fit model [1SE] separately for males and females and calculating the difference in predicted values (male − female) at representative PW values corresponding to the 25th, 50th, and 75th percentiles of PW (15.67, 17.13, and 18.46 mm, respectively) (Table 2). 95% confidence intervals were estimated from 500 bootstrap resamples^22^.

**Table 2.** Predicted male−female differences for the nine traits at representative PW values (25th, 50th, and 75th percentiles). The 25th, 50th, and 75th percentiles of PW were 15.67, 17.13, and 18.46 mm, respectively.

| Trait | Difference at PW25<br>[95% CI] | Difference at PW50<br>[95% CI] | Difference at PW75<br>[95% CI] |
| --- | --- | --- | --- |
| HL | 2.20 [2.09, 2.31] | 3.70 [3.60, 3.82] | 3.91 [3.80, 3.99] |
| PDL | 1.44 [1.38, 1.50] | 1.66 [1.59, 1.72] | 1.64 [1.58, 1.70] |
| PDW | 0.97 [0.87, 1.04] | 1.34 [1.28, 1.42] | 1.44 [1.38, 1.50] |
| BL | -0.25 [-0.36, -0.16] | -0.29 [-0.38, -0.20] | -0.32 [-0.45, -0.19] |
| EL | -0.23 [-0.30, -0.19] | -0.30 [-0.37, -0.25] | -0.45 [-0.50, -0.39] |
| ML | 0.04 [0.01, 0.01] | 0.02 [-0.18, 0.05] | 0.02 [-0.01, 0.06] |
| PL | 0.24 [0.20, 0.28] | 0.38 [0.35, 0.41] | 0.50 [0.46, 0.54] |
| AW | -0.22 [-0.26, -0.16] | -0.35 [-0.40, -0.30] | -0.40 [-0.46, -0.35] |
| AD | 0.02 [-0.04, 0.08] | -0.05 [-0.12, 0.01] | -0.10 [-0.16, -0.03] |

To test whether the observed nonlinearity could be explained by discrete dimorphism, we introduced two-component mixture regression (MIXTURE_K2) as an additional candidate model and compared it, within the same cross-validation framework, with the best of the eight continuous models, defined as the continuous model with the lowest nll. Specifically, we calculated Delta nll_dimorph = nll (best continuous model) − nll (MIXTURE_K2) (Table S4). We considered discrete dimorphism within the adult population to be supported only if Delta nll_dimorph was clearly positive and MIXTURE_K2 consistently outperformed the best continuous model.

Because HL and PDW emerged as the focal nonlinear traits, we evaluated the association between the horn and the pronotal depression in two complementary ways. First, we analyzed correlations among residuals obtained after removing the component explained by a GAM with PW as the predictor (Table 3; Fig. S7). Second, we fitted a GAM separately for each sex with PDW as the response and PW and HL as simultaneous predictors [PDW ∼ s(PW) + s(HL)] to assess whether body-size-independent covariation was present (Table 4; Fig. 4).

**Fig. 4.**
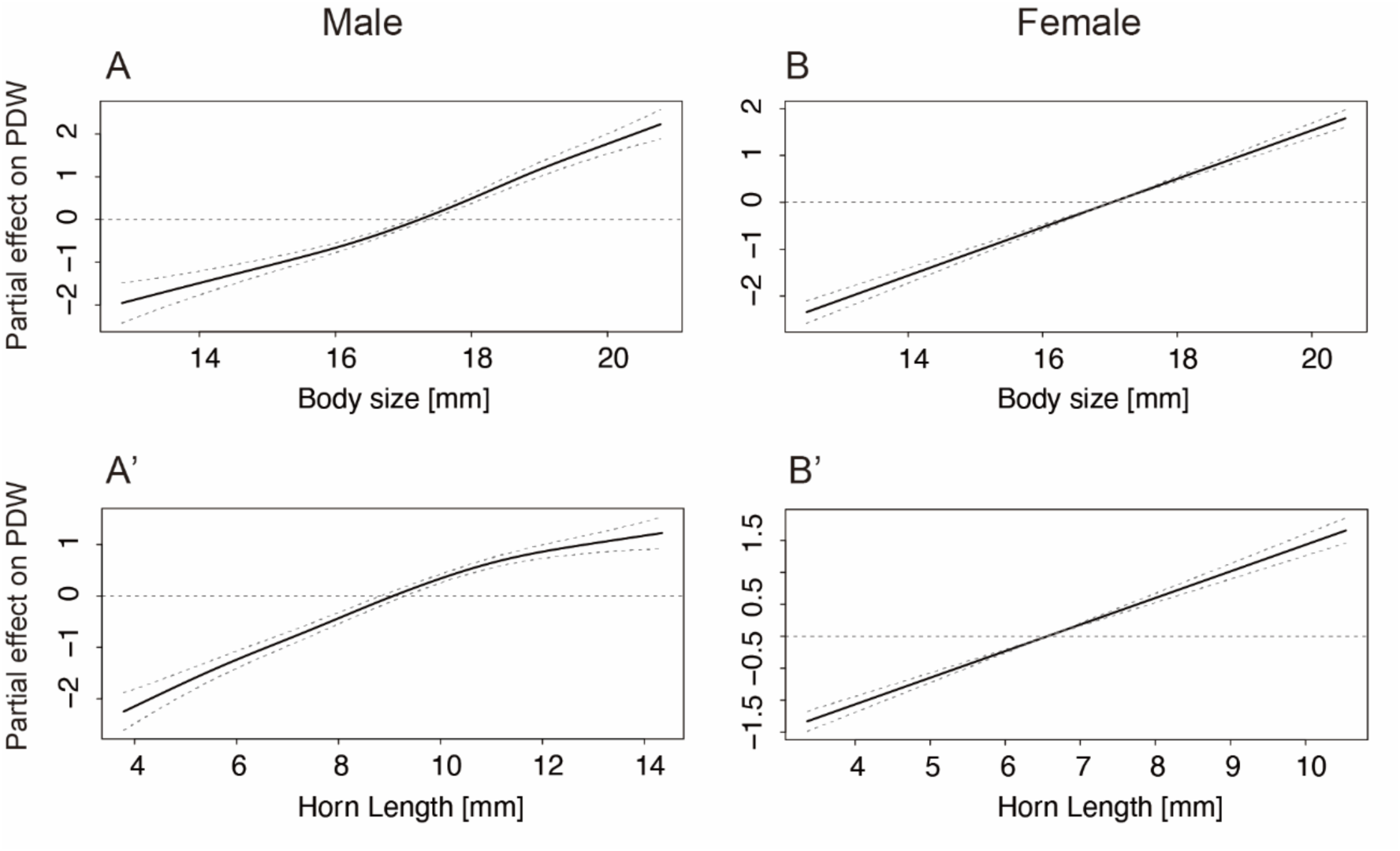
Partial effects of PW and HL in the GAM with PDW as the response variable. Partial effects from the GAM with PDW as the response and PW and HL as predictors. Each curve shows the estimated partial effect of one predictor while controlling for the other. Shown are the partial effects of PW (A) and HL (A′) in males, and of PW (B) and HL (B′) in females.

**Table 3.**
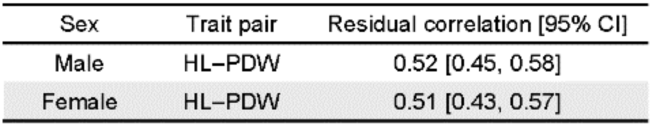
Pearson correlation between HL and PDW based on residuals after removing the component explained by a GAM with PW as the predictor.

**Table 4.**
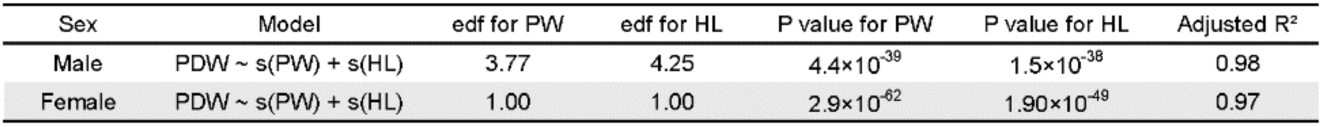
Summary of the GAM fitted to PDW as a function of PW and HL [PDW ∼ s(PW) + s(HL)]. edf = effective degrees of freedom

To evaluate robustness to the choice of body-size proxy, we repeated the key analyses using body length (BL) instead of PW. In these sensitivity analyses, HL, PDL, PDW, PW, EL, ML, PL, AW, and AD were analyzed relative to BL, and Best-fit model [1SE], Delta nll, Delta nll_HET, and Delta nll_dimorph were re-evaluated with the same candidate model set, cross-validation procedure, and one-standard-error rule described above. Sex differences in trait size were calculated at the 25th, 50th, and 75th percentiles of BL (38.44, 42.03, and 44.95 mm, respectively). We also repeated the residual-correlation analyses after removing the component explained by a GAM with BL as the predictor and fitted a GAM of the form PDW ∼ s(BL) + s(HL) to test whether the association between horn and pronotal depression was retained under the alternative body-size proxy (Tables S5-S9; Figs. S8-S9).

## Results

### Measurement reproducibility and influential observations

To assess reproducibility for PDL and PDW, the two measured traits most susceptible to differences in contour delineation, we remeasured 50 males and 50 females. ICC (A,1) ranged from 0.990 to 0.995, CCC from 0.989 to 0.995, and relative TEM from 1.13 to 1.80%, indicating high reproducibility for both traits (Table S1).

We also examined overlap in body-size (PW) distributions and the frequency of influential observations, because both can affect the stability of nonlinear estimation and sex comparisons. PW ranged from 12.86 to 20.78 mm in males and from 12.49 to 20.50 mm in females, providing broad overlap across the analytical range. Based on Cook’s distance and studentized residuals, 44 influential observations were identified across all nine traits (Table S2). Their frequency within each trait-by-sex group was nevertheless low, with at most 0–5 observations per group, and many appeared as isolated points in the scatterplots (Fig. S4). We therefore retained all observations in the subsequent analyses.

### Detecting nonlinearity by model comparison

For each trait, we asked whether static allometry was adequately described by a linear model or whether a continuous nonlinear model was better supported on the basis of generalization performance. Repeated cross-validation showed that Delta nll = nll (LINEAR_HOM) − nll (Best-fit model [1SE]) was relatively large and positive for HL and PDW in both sexes, indicating that continuous nonlinear models consistently improved predictive performance over the linear model for these traits (Fig. 3; Table 1). Specifically, Delta nll for HL was 0.186 in males and 0.387 in females, whereas Delta nll for PDW was 0.095 in males and 0.092 in females. Moderate improvement was also observed for male PDL (0.053) and male EL (0.044). For most other trait-by-sex combinations, improvements were generally small, with Delta nll values often remaining around 0.001–0.020 (Fig. S6; Table 1).

For groups in which the Best-fit model [1SE] was a HET model, we additionally calculated Delta nll_HET = nll (LINEAR_HET) − nll (Best-fit model [1SE]). For HL (0.197 in males and 0.395 in females) and PDW (0.105 in males and 0.094 in females), Delta nll_HET closely matched Delta nll (Table 1), indicating that the improvement was attributable mainly to nonlinearity in the mean structure rather than to heteroscedasticity alone. By contrast, for male EL, which showed moderate improvement overall, Delta nll_HET decreased to 0.024, suggesting a larger contribution of variance heterogeneity, including increased residual variance at larger body sizes (Table 1). Taken together, these results indicate strong mean-structure nonlinearity for HL and PDW, with weaker and more sex-specific evidence for EL.

### Best-fit models for HL and PDW, and sex differences in predicted values

For HL and PDW, the two traits that showed the strongest nonlinearity, we fitted the Best-fit model [1SE] to the full dataset and examined the fitted curves together with their 95% confidence bands (Fig. 2). For HL, LOGISTIC_HET was selected for males, whereas GAM_HET was selected for females, indicating that a more flexible curve was required for females (Fig. 2; Table 1). For PDW, LOGISTIC_HET was selected for males and SEGMENTED_HET for females, suggesting a female-specific change in slope over a restricted PW range (Fig. 2; Table 1). Fitted curves and 95% confidence bands for the other seven traits are shown in Fig. S5.

These results indicate that the Best-fit model [1SE] for HL and PDW differed between males and females. We therefore quantified sex differences by estimating predicted male-female differences at representative PW values (25th, 50th, and 75th percentiles; PW = 15.67, 17.13, and 18.46 mm). For both HL and PDW, predicted values were larger in males at all three points, and the magnitude of the difference increased in the medium-to-large PW range (HL: 2.20, 3.70, and 3.91 mm; PDW: 0.97, 1.34, and 1.44 mm) (Table 2). Thus, sexual dimorphism in HL and PDW was evident both in fitted curve shape and in predicted values at representative PW values.

### Mixture-model convergence and the discrete-dimorphism hypothesis

To test whether the nonlinear scaling observed in HL and PDW within each sex could be explained as discrete dimorphism, we additionally fitted two-component mixture regression (MIXTURE_K2). MIXTURE_K2 generally converged satisfactorily under cross-validation, allowing comparison on the same nll scale (Table S4). However, MIXTURE_K2 did not outperform the best continuous model for either HL or PDW. The resulting Delta nll_dimorph = nll (best continuous model) − nll (MIXTURE_K2) was −0.18 for males and −0.31 for females in HL, and −0.10 for males and −0.09 for females in PDW (Table S4). Thus, at least within the scope of the present data and candidate model set, we found no positive evidence for discrete within-sex dimorphism in either HL or PDW.

### Covariation after accounting for body size

To determine whether the horn and pronotal depression covaried beyond their shared dependence on body size, we analyzed their association after adjusting for PW. Residual correlations after removing the component explained by a GAM with PW as the predictor showed a moderate positive association between HL and PDW in both males (r = 0.52) and females (r = 0.51) (Table 3; Fig. S7). In addition, in GAMs with PDW as the response and both PW and HL as predictors [PDW ∼ s(PW) + s(HL)], HL contributed additional structure in both sexes (Table 4; Fig. 4). The shape of the fitted relationship differed between the sexes: in males, both smooth terms were clearly nonlinear (edf = 3.77 for PW and 4.25 for HL), whereas in females, both edf values were close to 1, indicating an approximately linear relationship. Because the high adjusted R^2 values (0.97–0.98; Table 4) probably reflect the dominant contribution of PW rather than the effect of HL alone, we interpret this pattern conservatively as size-independent covariation rather than as direct evidence of shared developmental control.

### Sensitivity analyses using BL as the body-size proxy

Because the primary body-size proxy, PW, is derived from the pronotum, we assessed the robustness of the main conclusions by repeating the key analyses with BL as an alternative proxy. HL again showed the largest Delta nll values in both sexes (0.209 in males and 0.314 in females), and PDW also retained relatively large improvements (0.105 in males and 0.066 in females) (Table S5). For both HL and PDW, predicted values remained larger in males across representative BL values, and the magnitude of the sex difference again increased in the medium-to-large size range (Table S6). These patterns were therefore broadly consistent with those obtained in the primary PW-based analyses.

Likewise, MIXTURE_K2 did not outperform the best continuous model for either HL or PDW in either sex, and all Delta nll_dimorph values were negative (Table S7).

Residual correlations after removing the component explained by a GAM with BL as the predictor again showed a positive association between HL and PDW, with r = 0.65 in males and r = 0.60 in females (Table S8). The overall correlation structure among all traits based on BL-adjusted residuals is shown in Fig. S9. In addition, the GAM of the form PDW ∼ s(BL) + s(HL) retained positive partial effects of both BL and HL in both sexes, although the complexity of the smooth terms differed from that in the primary analyses. In males, the edf values for both terms were close to 1, indicating an approximately linear relationship, whereas in females more flexible curves were required (edf = 1.78 for the BL term and 3.51 for the HL term) (Table S9; Fig. S8). Thus, the positive association between the horn and the pronotal depression was robust to body-size proxy choice, even though the detailed shape of the fitted smooths depended to some extent on the proxy used.

## Discussion

Using a large morphological dataset of 500 males and 500 females, we (i) compared the static allometry of nine traits relative to PW within a candidate set that treated mean structure and variance structure symmetrically, (ii) quantified sexual dimorphism from the best-fit models and from predicted values at representative PW values, (iii) tested whether the observed nonlinearity could be explained as discrete dimorphism by comparing MIXTURE_K2 with continuous nonlinear models, (iv) evaluated covariation between the horn and the pronotal depression after accounting for body size, and (v) assessed the robustness of the main conclusions by repeating the analyses with BL. A notable feature of our approach is that apparent nonlinear improvement could be partitioned into contributions from mean structure and variance structure. This made it possible to distinguish cases in which a better fit reflected genuine nonlinearity in the mean from cases in which it primarily reflected heteroscedasticity. In addition, by comparing MIXTURE_K2 and continuous nonlinear models on the same nll scale, we provide a predictive framework for evaluating discrete dimorphism against continuous alternatives within the tested model set.

### Nonlinear scaling across traits

Within a common cross-validation framework, continuous nonlinear models consistently outperformed the linear homoscedastic model for HL and PDW in both sexes, supporting clear nonlinearity in the static allometry of both the exaggerated trait and the related pronotal trait. In contrast, many other traits were adequately approximated by linear models, indicating that traits with stronger and weaker nonlinearity coexist within the same population. HET models were selected for some traits, showing that residual variance can change with body size. However, because the contributions of mean structure and variance structure were evaluated separately, the substantial improvement for HL and PDW is unlikely to be explained solely by heteroscedasticity and instead most plausibly reflects nonlinearity in the mean structure itself. These trait-specific differences suggest that size-related scaling is not uniform across the body plan and may reflect trait-specific growth dynamics.

### Sexual dimorphism in HL and PDW

For several traits, the Best-fit model [1SE] selected based on generalization performance differed between the sexes (Table 1; Fig. 2; Fig. S5). This contrast was most evident for HL and PDW, for which the best-fit mean structures differed between males and females. Predicted values at representative PW values were also consistently larger in males for both traits, and the magnitude of the difference increased in the medium-to-large size range (Table 2). Sexual dimorphism in HL and PDW was therefore evident not only in fitted curve shape but also in predicted trait values at representative body sizes. By contrast, the other six traits showed no comparably strong sex differences (BL, EL, ML, PL, AW, and AD) (Table 2). Overall, sexual dimorphism was most pronounced in HL and PDW, consistent with sex-specific scaling relationships for these traits.

### Continuous nonlinearity versus discrete dimorphism

Whether nonlinear scaling reflects discrete dimorphism within an adult population is a central question in the study of exaggerated traits^5^. Here we compared the best continuous model and MIXTURE_K2 separately within each sex under the same cross-validation framework. For neither HL nor PDW, despite their clear nonlinearity, did MIXTURE_K2 outperform the best continuous model. Within the scope of the present data and candidate model set, these results indicate that the observed patterns in HL and PDW can be explained without invoking discrete within-sex dimorphism.

Discrete horn dimorphism is well documented in some horned beetles. In *Onthophagus taurus*, for example, males above a body-size threshold develop long horns, whereas smaller males are short-horned or effectively hornless, and these morphs are associated with alternative reproductive tactics^23,24^. By contrast, although HL in *O. rhinoceros* showed clear nonlinearity, our model comparisons did not provide positive support for discrete within-sex dimorphism; the observed pattern was adequately captured by a continuous nonlinear model. The head horn has been reported to assist adults in penetrating palm tissue^25^, suggesting a functional role in boring behavior. Nevertheless, our results do not rule out other latent mixture structures or developmental thresholds that were not captured by the present model set. The functional significance of horn and pronotal depression variation therefore remains to be tested directly through behavioral and developmental studies.

### Size-independent covariation between the horn and the pronotal depression

In horned beetles, the head and pronotum are the principal regions of horn formation^7^. In *O. rhinoceros*, both residual-correlation analyses and partial-regression analyses showed that HL and PDW remained positively associated after adjusting for body size. This suggests that their covariation cannot be explained solely by overall size. Importantly, the association was retained not only in the primary PW-based analyses but also in the BL sensitivity analyses. The pattern is therefore difficult to dismiss as a simple artefact of using PW or of the fact that PW and PDW are both pronotal measurements. This strengthens the inference that the horn-pronotal association is biologically meaningful.

### Interpreting nonlinear scaling in exaggerated traits

Exaggerated traits in horned beetles are often discussed in terms of thresholds and discrete morphs^23,24,26^. A central contribution of this study is to show that pronounced nonlinearity in adult static allometry does not, by itself, justify a dimorphism interpretation. By comparing continuous nonlinear models and mixture models on a common predictive scale, we provide a stricter basis for distinguishing between these alternatives. More broadly, by analyzing the horn, the pronotal depression, and several additional traits within the same framework, we show that strong nonlinearity is trait-specific rather than a global property of body-size scaling in this species. This comparative perspective should be useful for future studies seeking to connect adult morphology with developmental and functional mechanisms.

## Conclusion

In *O. rhinoceros*, adult horn length and pronotal depression width show continuous nonlinear static allometry relative to body size. In the primary PW-based analyses, clear nonlinearity was detected in HL and PDW, and these main conclusions were broadly retained in the BL sensitivity analyses. Within the scope of the present data and candidate model set, two-component mixture regression did not outperform the best continuous model in either sex, providing no positive support for discrete within-sex dimorphism. In addition, the positive association between the horn and the pronotal depression persisted after adjustment for both PW and BL. Thus, the association itself was robust to body-size proxy choice, even though the detailed smooth shapes depended somewhat on the proxy used. Together, these results provide a quantitative foundation for future experimental work on the developmental mechanisms and functional significance of exaggerated and related traits in this species.

## Data availability

All data generated or analyzed during this study are included in this published article and its Supporting Information files.

## Acknowledgements

We thank Shigeru Okiyama (Minami-Daito Village Office) and Kazuhiro Kikuchi (Minami-Daito Village Agricultural Youth Club) for providing specimens, the Model Plant Research Facility/NIBB BioResource Center, the Emerging Model Organisms Facility/NIBB Trans-Scale Biology Center, and Toshiyuki Sazi of the Section of Instrument Design Room at NIPS. This work was supported by JSPS KAKENHI Grant Numbers 21K15135 and 25K02028 (to S. M.), the Sumitomo Foundation Grant Number 2402226 (to S. M.), and MEXT KAKENHI Grant Number 20H05944 (to T. N.).

## Author contributions

M. O.: Data curation, Validation, Visualization, Formal analysis, Writing – original draft; T. N.: Writing – original draft, Funding acquisition, Supervision.; S. M.: Conceptualization, Data curation, Methodology, Validation, Visualization, Formal analysis, Writing – original draft, Funding acquisition, Project administration, Supervision.

## Supporting information

**Fig. S1.**
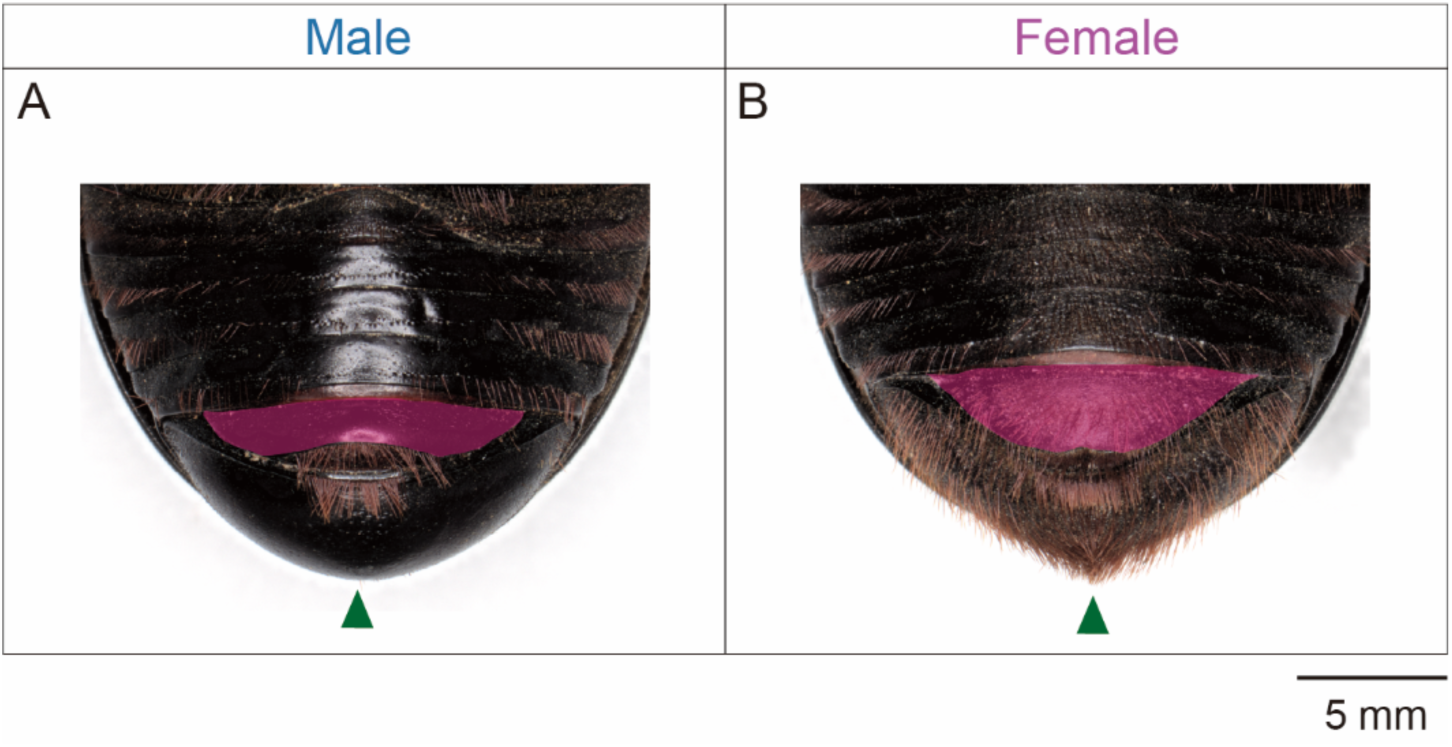
Sexing of adult *O. rhinoceros* based on abdominal morphology. The magenta outline indicates the terminal abdominal sternite, and the green arrowheads indicate terminal abdominal setae. (A) Ventral abdomen of a male. The terminal abdominal sternite is smooth, deeply emarginate posteriorly, and the pygidium is bare. (B) Ventral abdomen of a female. The terminal abdominal sternite bears fine setae and is less deeply emarginate posteriorly than in the male. Fine setae are also present on the pygidium.

**Fig. S2.**
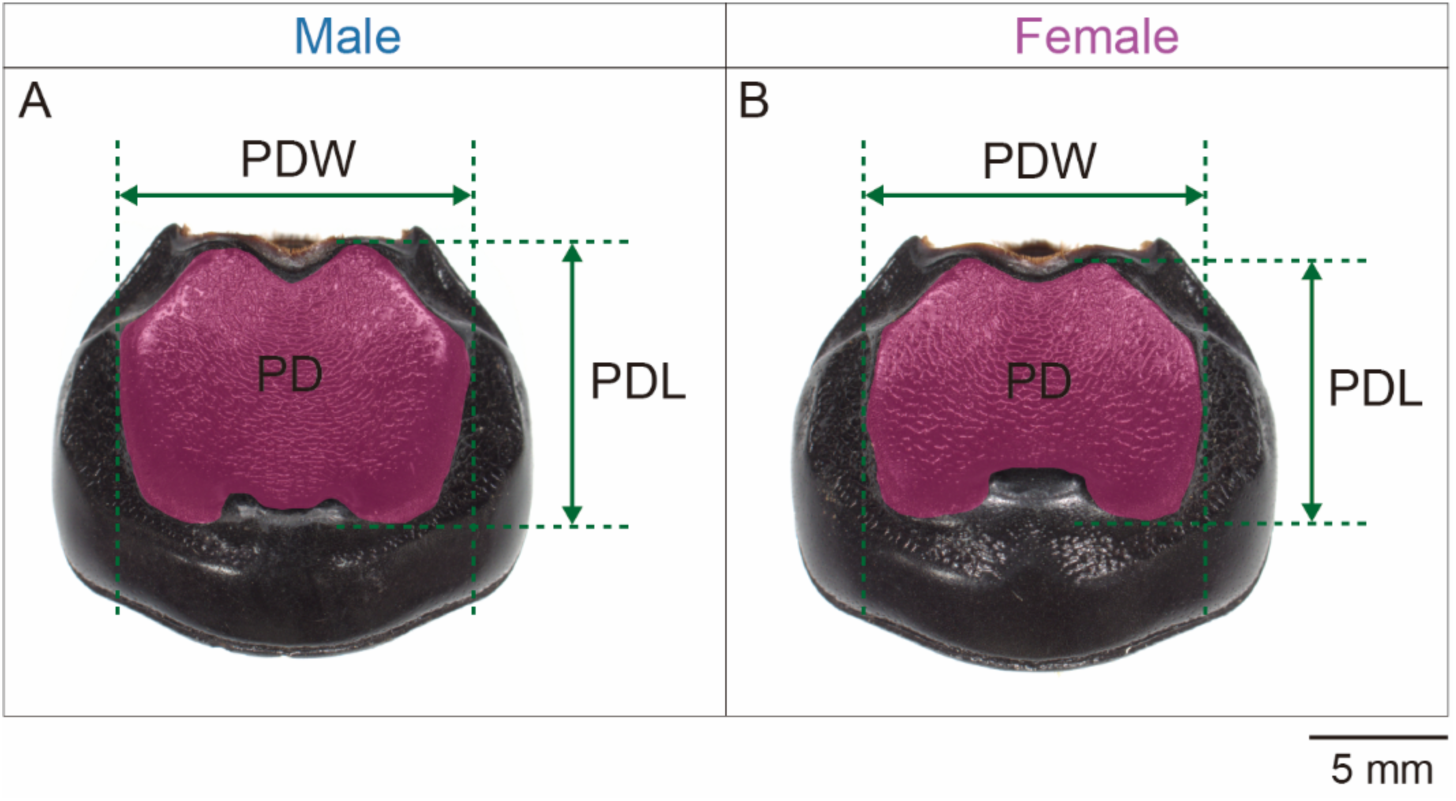
Definition of the pronotal depression and measurement sites. Pronotum of an adult male (A) and an adult female (B) of *O. rhinoceros*. The magenta-outlined region was defined as the pronotal depression (PD). Its maximum length and maximum width were measured as PDL and PDW, respectively.

**Fig. S3.**
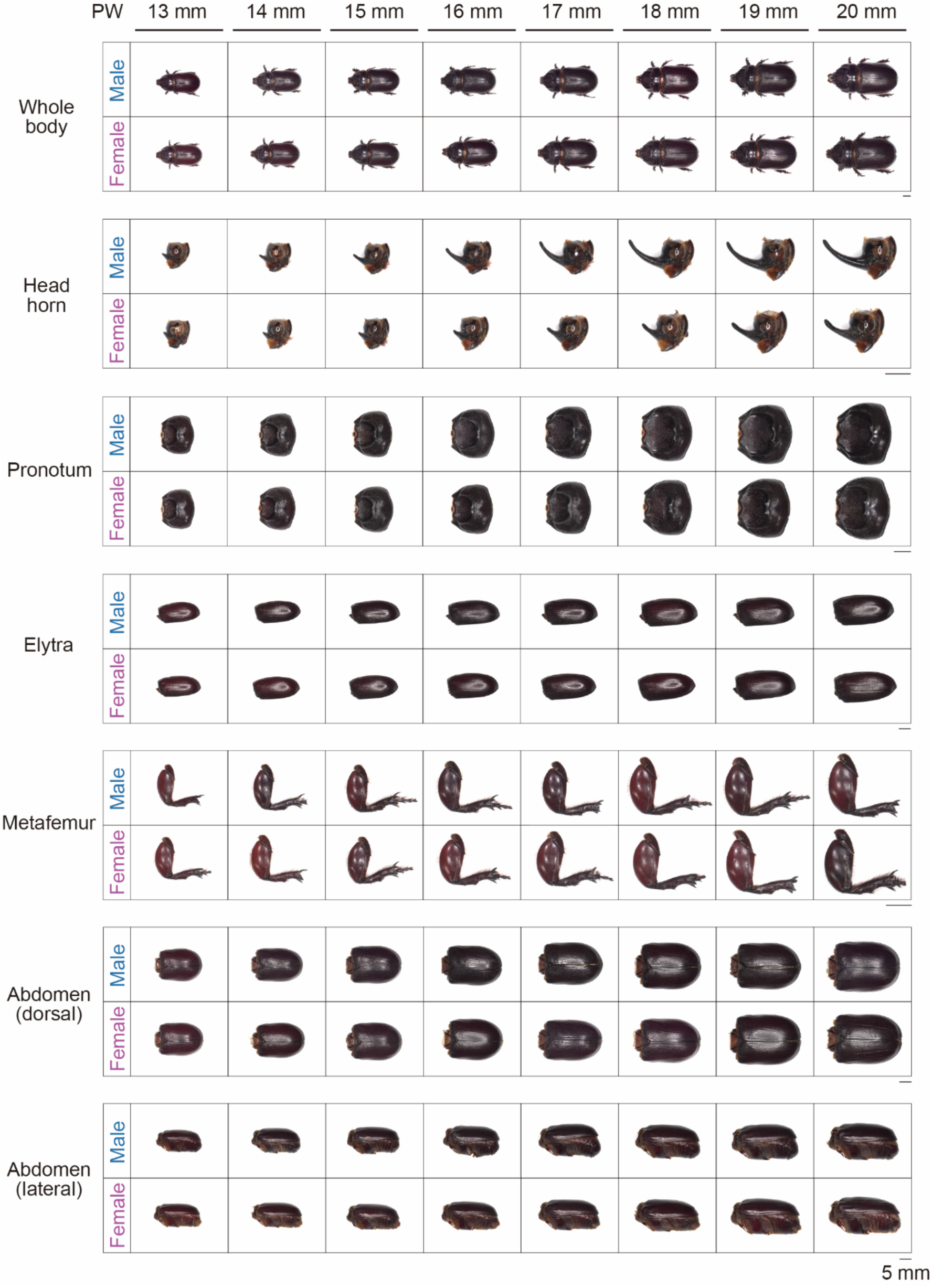
Comparison of male and female traits at representative PW values. Representative males and females were selected at 1-mm PW intervals from 13.0 to 20.0 mm to compare trait morphology between the sexes. For each trait, the anterior side is oriented to the left and the posterior side to the right. Some individuals lacked tarsal segments or other appendages, but such damage did not affect measurement of the nine traits analyzed in this study.

**Fig. S4.**
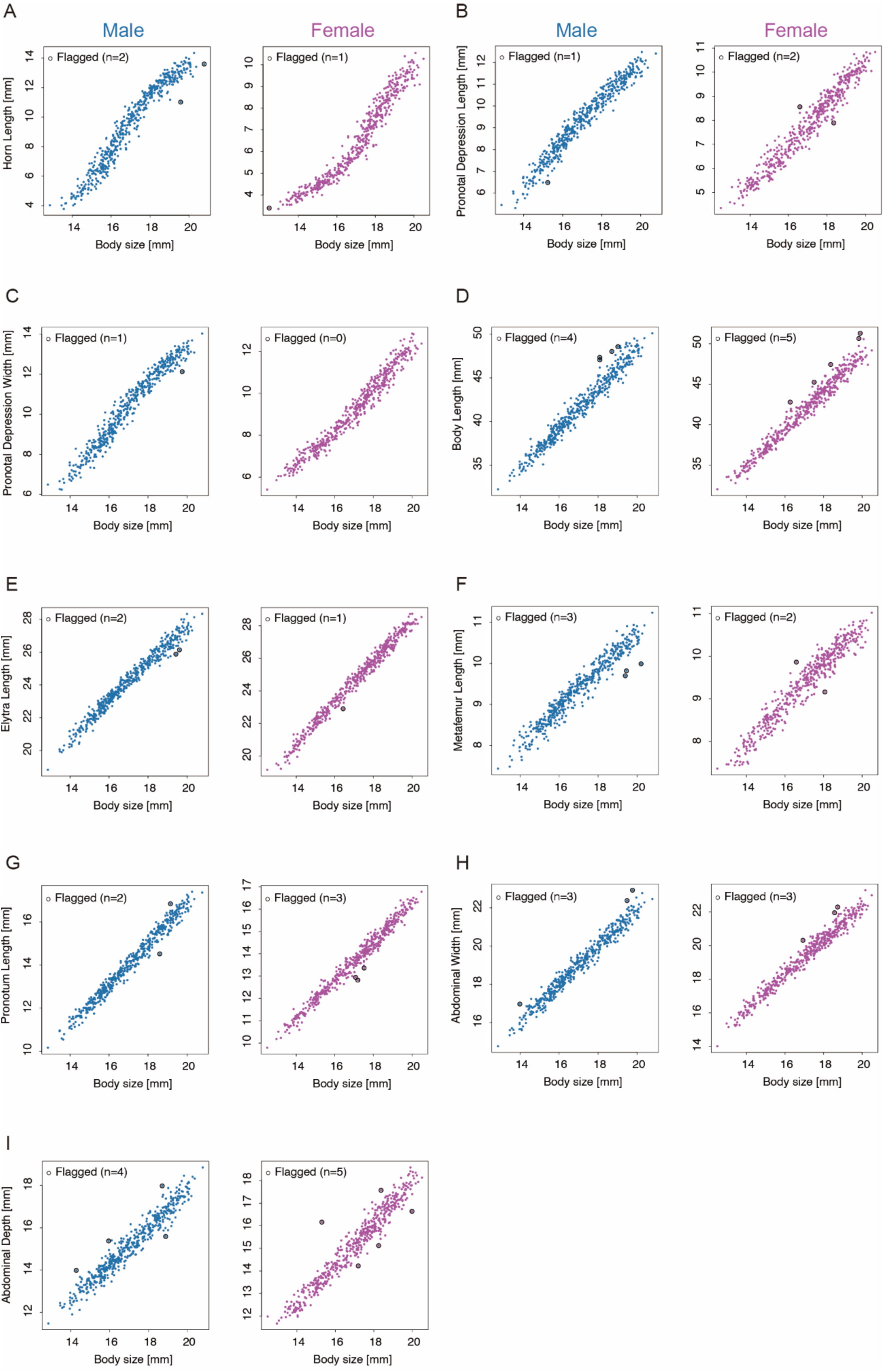
Scatterplots of all nine traits and visualization of influential points. Scatterplots show trait values against PW for each group. Influential observations identified by Cook’s distance and studentized residuals are outlined in black. Males are shown in blue and females in magenta. (A) HL, (B) PDL, (C) PDW, (D) BL, (E) EL, (F) ML, (G) PL, (H) AW, and (I) AD.

**Fig. S5.**
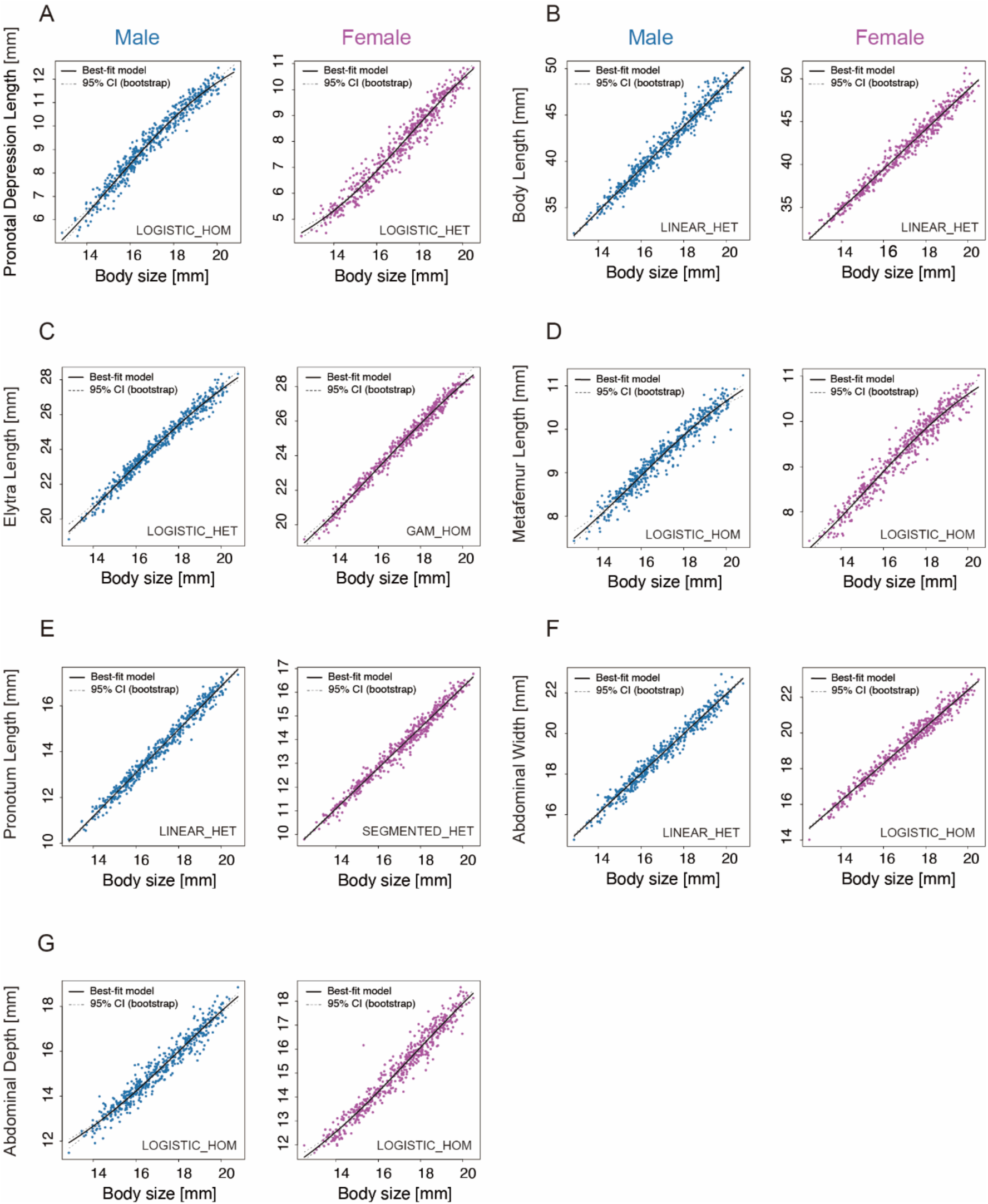
Estimated static allometry curves and 95% confidence bands for the seven traits other than HL and PDW. Fitted curves for the seven remaining traits against PW are shown with the 95% confidence bands. Best-fit model [1SE] is shown by the solid black line, and the 95% confidence bands are shown by gray dashed lines. Males are shown in blue and females in magenta. (A) PDL, (B) BL, (C) EL, (D) ML, (E) PL, (F) AW, and (G) AD.

**Fig. S6.**
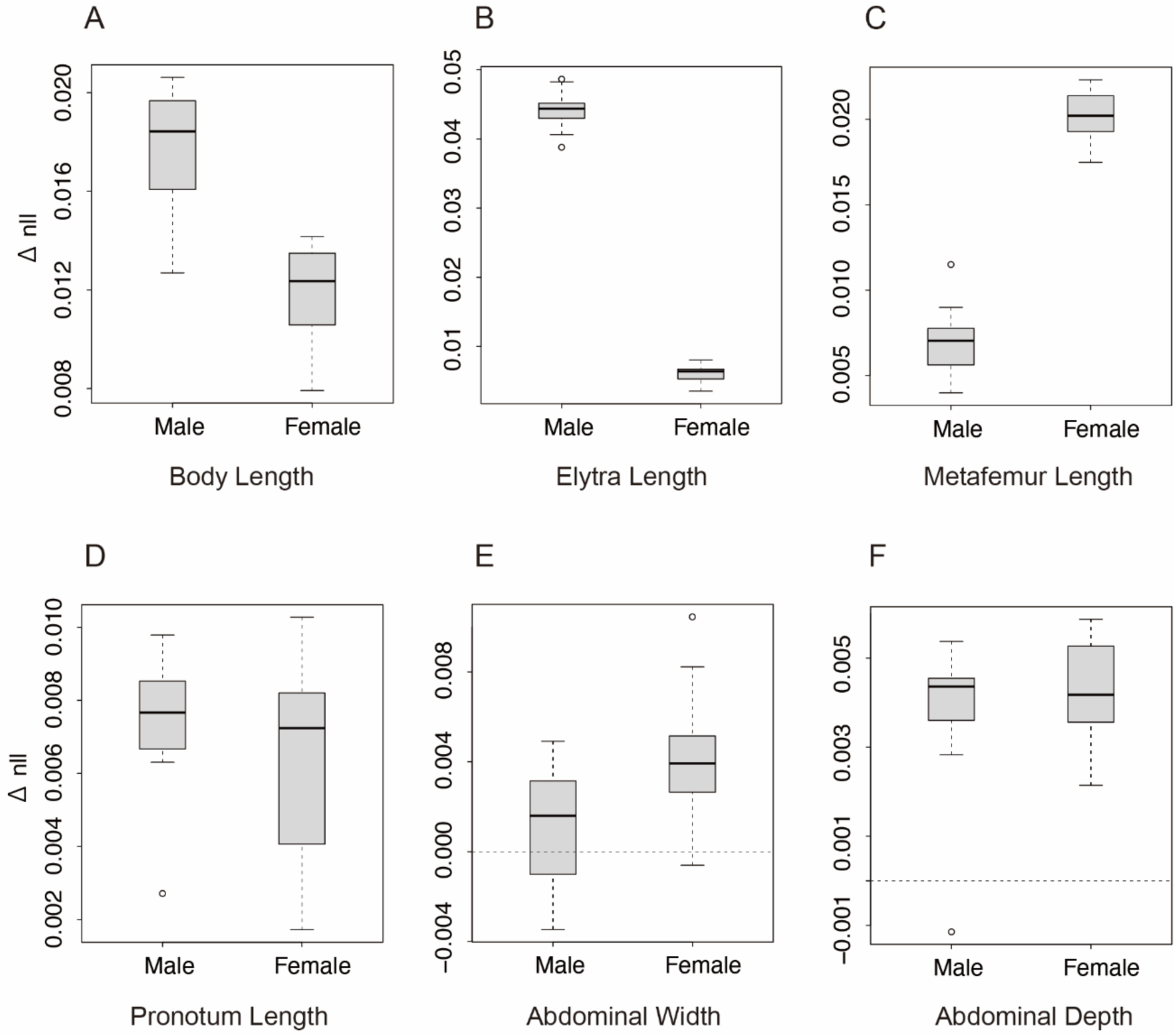
Distribution of Δnll from repeated cross-validation for the six traits other than HL, PDL, and PDW. For each trait, boxplots show Δnll values obtained from repeated cross-validation (20 repeats). Open circles indicate outliers. (A) BL, (B) EL, (C) ML, (D) PL, (E) AW, and (F) AD.

**Fig. S7.**
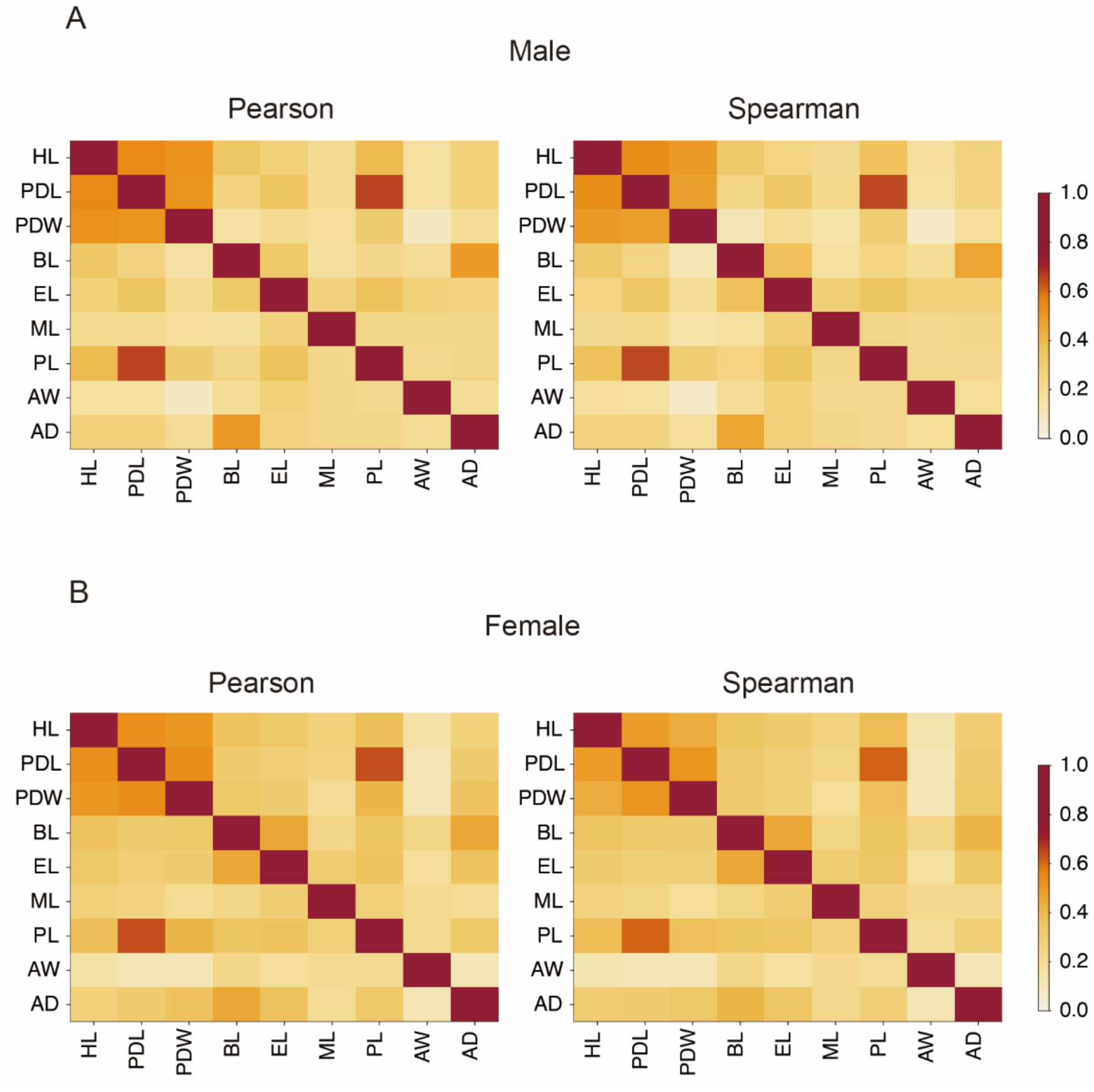
Heat maps of residual correlations after removing the effect of PW. Residuals were obtained after removing the component explained by a GAM with PW as the predictor. Correlations among the nine traits are shown as matrices. Pearson correlations are shown on the left and Spearman rank correlations on the right. Color intensity indicates the correlation coefficient (0–1), and diagonal cells show self-correlation (r = 1). (A) Males. (B) Females.

**Fig. S8.**
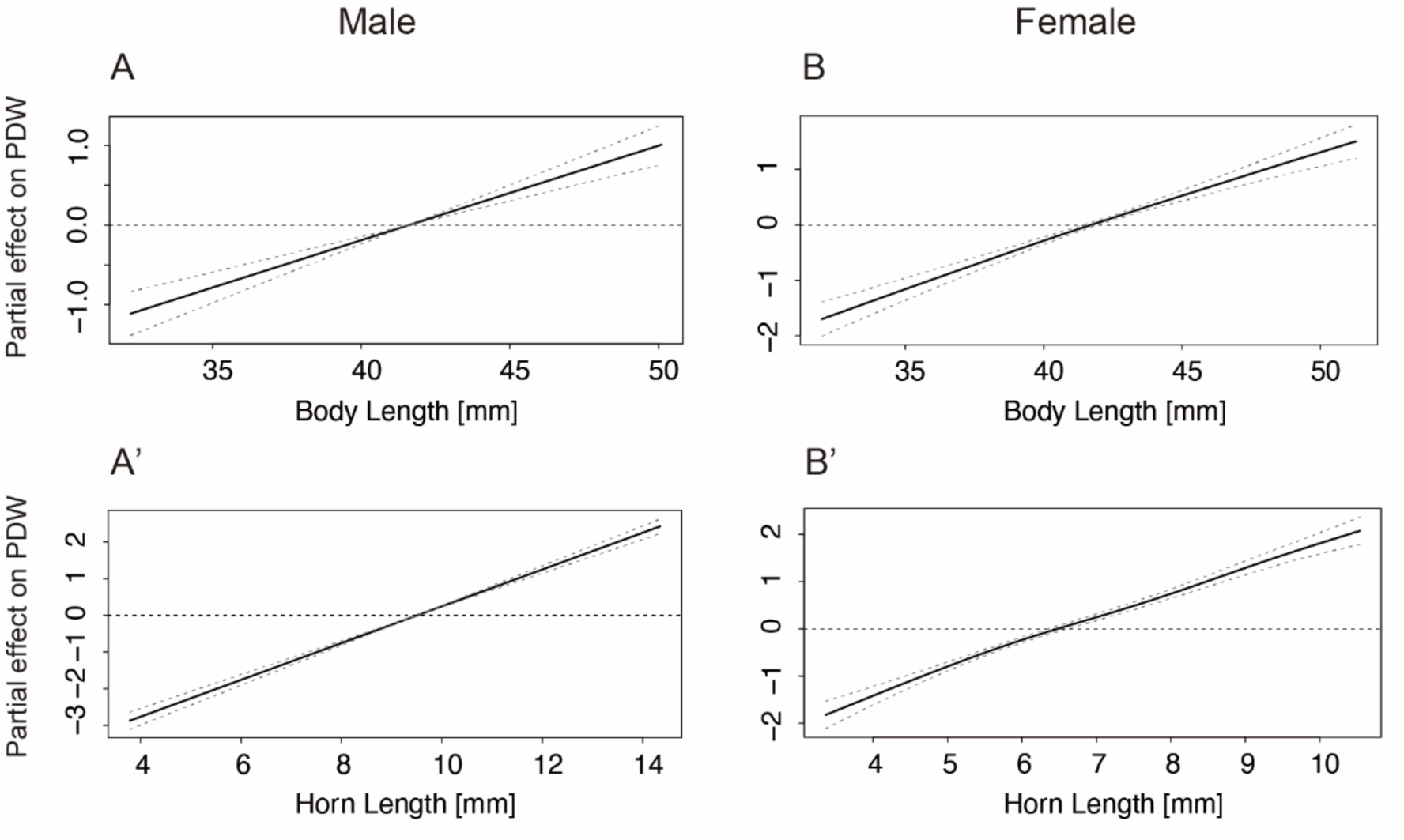
Partial effects of BL and HL in the GAM with PDW as the response variable. Partial effects from the GAM with PDW as the response and BL and HL as predictors. Shown are the partial effects of BL (A) and HL (A′) in males, and of BL (B) and HL (B′) in females.

**Fig. S9.**
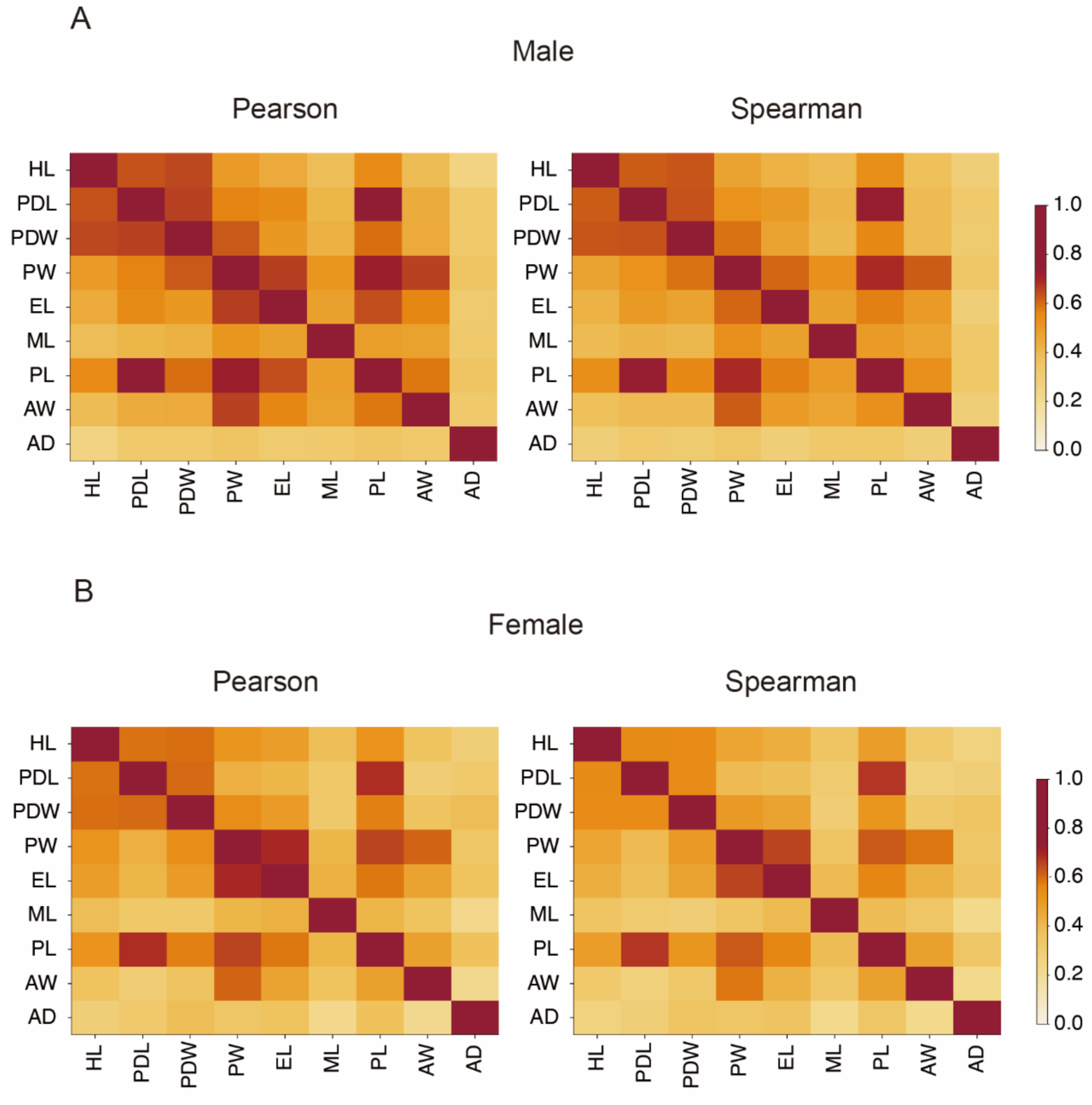
Heat maps of residual correlations after removing the effect of BL. Residuals were obtained after removing the component explained by a GAM with BL as the predictor. Correlations among the nine traits are shown as matrices. Pearson correlations are shown on the left and Spearman rank correlations on the right. Color intensity indicates the correlation coefficient (0–1), and diagonal cells show self-correlation (r = 1). (A) Males. (B) Females.

**Table S1.**
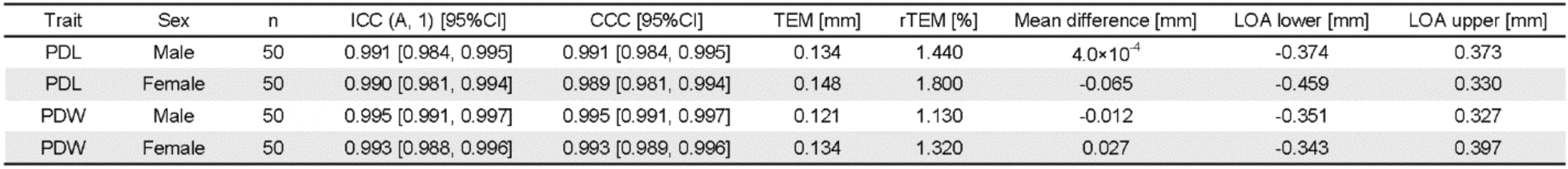
Measurement reproducibility for PDL and PDW. rTEM = relative TEM; LOA = limits of agreement.

**Table S2.** Number of influential observations identified by Cook’s distance and studentized residuals.

| Trait | Male | Female |
| --- | --- | --- |
| HL | 2 | 1 |
| PDL | 1 | 2 |
| PDW | 1 | 0 |
| BL | 4 | 5 |
| EL | 2 | 1 |
| ML | 3 | 2 |
| PL | 2 | 3 |
| AW | 3 | 3 |
| AD | 4 | 5 |

**Table S3.** Estimated parameters of Best-fit model [1SE] for HL and PDW. x_0_ = inflection point; edf = effective degrees of freedom; psi = breakpoint; Slope1 = slope before the breakpoint; Slope2 = slope after the breakpoint.

| Trait | Sex | Best-fit model [1SE] | $x_0$ [95%CI] | k [95%CI] | edf [95%CI] | psi [95%CI] | Slope1 [95%CI] | Slope2 [95%CI] |
| --- | --- | --- | --- | --- | --- | --- | --- | --- |
| HL | Male | LOGISTIC_HET | 16.41 [16.29, 16.52] | 0.76 [0.68, 0.83] | — | — | — | — |
| PDW | Male | LOGISTIC_HET | 16.43 [16.22, 16.62] | 0.57 [0.49, 0.65] | — | — | — | — |
| HL | Female | GAM_HET | — | — | 7.59 [7.24, 8.44] | — | — | — |
| PDW | Female | SEGMENTED_HET | — | — | — | 15.74 [15.51, 16.40] | 0.69 [0.65, 0.77] | 1.03 [1.00, 1.07] |

**Table S4.**
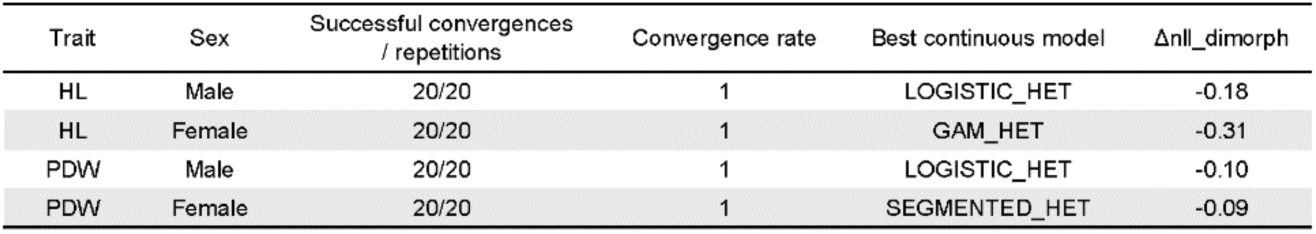
Convergence of mixture regression (MIXTURE_K2) and Δnll_dimorph for HL and PDW across repeated cross-validation (20 repeats). Δnll_dimorph = nll (best continuous model) − nll (MIXTURE_K2).

| Trait | Sex | Successful convergences / repetitions | Convergence rate | Best continuous model | $\Delta nll\_dimorph$ |
| --- | --- | --- | --- | --- | --- |
| HL | Male | 20/20 | 1 | LOGISTIC_HET | -0.18 |
| HL | Female | 20/20 | 1 | GAM_HET | -0.31 |
| PDW | Male | 20/20 | 1 | LOGISTIC_HET | -0.10 |
| PDW | Female | 20/20 | 1 | SEGMENTED_HET | -0.09 |

**Table S5.**
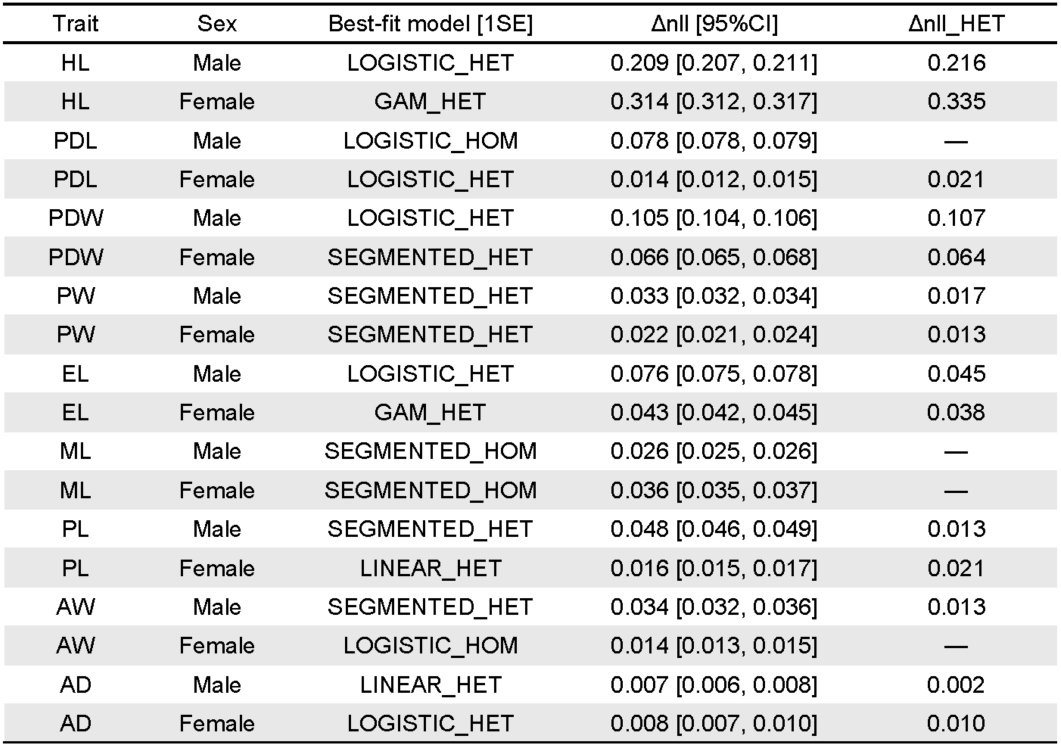
Best-fit model [1SE] and the associated Δnll and Δnll_HET values from repeated cross-validation using BL as the body-size proxy.

| Trait | Sex | Best-fit model [1SE] | $\Delta nll$ [95%CI] | $\Delta nll\_HET$ |
| --- | --- | --- | --- | --- |
| HL | Male | LOGISTIC_HET | 0.209 [0.207, 0.211] | 0.216 |
| HL | Female | GAM_HET | 0.314 [0.312, 0.317] | 0.335 |
| PDL | Male | LOGISTIC_HOM | 0.078 [0.078, 0.079] | — |
| PDL | Female | LOGISTIC_HET | 0.014 [0.012, 0.015] | 0.021 |
| PDW | Male | LOGISTIC_HET | 0.105 [0.104, 0.106] | 0.107 |
| PDW | Female | SEGMENTED_HET | 0.066 [0.065, 0.068] | 0.064 |
| PW | Male | SEGMENTED_HET | 0.033 [0.032, 0.034] | 0.017 |
| PW | Female | SEGMENTED_HET | 0.022 [0.021, 0.024] | 0.013 |
| EL | Male | LOGISTIC_HET | 0.076 [0.075, 0.078] | 0.045 |
| EL | Female | GAM_HET | 0.043 [0.042, 0.045] | 0.038 |
| ML | Male | SEGMENTED_HOM | 0.026 [0.025, 0.026] | — |
| ML | Female | SEGMENTED_HOM | 0.036 [0.035, 0.037] | — |
| PL | Male | SEGMENTED_HET | 0.048 [0.046, 0.049] | 0.013 |
| PL | Female | LINEAR_HET | 0.016 [0.015, 0.017] | 0.021 |
| AW | Male | SEGMENTED_HET | 0.034 [0.032, 0.036] | 0.013 |
| AW | Female | LOGISTIC_HOM | 0.014 [0.013, 0.015] | — |
| AD | Male | LINEAR_HET | 0.007 [0.006, 0.008] | 0.002 |
| AD | Female | LOGISTIC_HET | 0.008 [0.007, 0.010] | 0.010 |

**Table S6.** Predicted male− female differences for HL and PDW at representative BL values (25th, 50th, and 75th percentiles). The 25th, 50th, and 75th percentiles of BL were 38.44, 42.03, and 44.95 mm, respectively.

| Trait | Difference at BL25<br>[95% CI] | Difference at BL50<br>[95% CI] | Difference at BL75<br>[95% CI] |
| --- | --- | --- | --- |
| HL | 2.29 [2.18, 2.41] | 3.94 [3.84, 4.07] | 4.09 [3.95, 4.20] |
| PDW | 1.05 [0.93, 1.12] | 1.53 [1.45, 1.62] | 1.61 [1.52, 1.69] |

**Table S7.** Convergence of MIXTURE_K2 and Δnll_dimorph when BL is used as the body-size proxy.

| Trait | Sex | Successful convergences<br>/ repetitions | Convergence rate | Best continuous model | $\Delta nll\_dimorph$ |
| --- | --- | --- | --- | --- | --- |
| HL | Male | 20/20 | 1 | LOGISTIC_HET | -0.22 |
| HL | Female | 20/20 | 1 | GAM_HET | -0.26 |
| PDW | Male | 20/20 | 1 | LOGISTIC_HET | -0.10 |
| PDW | Female | 20/20 | 1 | GAM_HET | -0.06 |

**Table S8.**
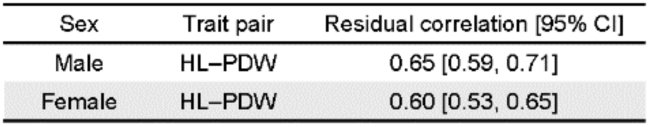
Pearson correlation between HL and PDW based on residuals after removing the component explained by a GAM with BL as the predictor.

**Table S9.**
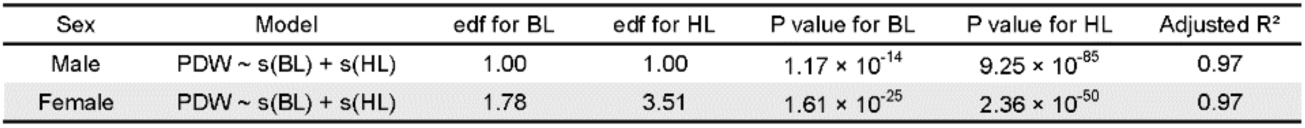
Summary of the GAM fitted to PDW as a function of BL and HL [PDW ∼ s(BL) + s(HL)].

